# An optimized toolkit for precision base editing

**DOI:** 10.1101/303131

**Authors:** Maria Paz Zafra, Emma M Schatoff, Alyna Katti, Miguel Foronda, Marco Breinig, Anabel Y. Schweitzer, Amber Simon, Teng Han, Sukanya Goswami, Emma Montgomery, Jordana Thibado, Francisco J. Sánchez-Rivera, Junwei Shi, Christopher R Vakoc, Scott W Lowe, Darjus F. Tschaharganeh, Lukas E Dow

**Affiliations:** Sandra and Edward Meyer Cancer Center, Department of Medicine, Weill Cornell Medicine, New York, NY; Weill Cornell / Rockefeller / Sloan Kettering Tri-I MD-PhD program, New York, NY; Weill Cornell Graduate School of Medical Sciences, Weill Cornell Medicine, New York, NY; Helmholtz-University Group “Cell Plasticity and Epigenetic Remodeling”, German Cancer Research Center (DKFZ) & Institute of Pathology University Hospital, 69120 Heidelberg, Germany; Cancer Biology and Genetics, Memorial Sloan Kettering Cancer Center, New York, NY; Cold Spring Harbor Laboratory, New York, NY; Department of Cancer Biology, Perelman School of Medicine, University of Pennsylvania, PA; Howard Hughes Medical Institute, Memorial Sloan Kettering Cancer Center, New York, NY; Department of Biochemistry, Weill Cornell Medicine, New York, NY

**Author notes:** Equal contribution (Maria Paz Zafra, Emma M Schatoff). Correspondence to Lukas Dow.

**Keywords:** Base editing, BE3, CRISPR, APOBEC

## Abstract

CRISPR base editing is a potentially powerful technology that enables the creation of genetic mutations with single base pair resolution. By re-engineering both DNA and protein sequences, we developed a collection of constitutive and inducible base editing vector systems that dramatically improve the ease and efficiency by which single nucleotide variants can be created. This new toolkit is effective in a wide range of model systems, and provides a means for efficient *in vivo* somatic base editing.

---

Cas9-guided DNA base editing is an exciting tool for precise genetic modification. Base editors are hybrid proteins that tether DNA modifying enzymes to nuclease defective Cas9 variants. This enables the direct conversion of cytosine (C) to other bases (T, A, or G) ^1–4^, or adenine (A) to inosine/guanine (I/G) nucleic acids ^5,6^, enabling the creation or repair of disease-associated single nucleotide variants (SNVs), or molecular recording^7^. The BE3 base editor carries a rat APOBEC cytidine deaminase at the N-terminus of Cas9n (Cas9^D10A^) and a uracil glycosylase inhibitor (UGI) domain at the C-terminus. This construct has been shown to drive targeted C>T transitions at nucleotide positions 3-8 of the protospacer (Figure 1a) following transfection of either plasmid DNA, or pre-assembled ribonuclear particles (RNPs) ^8,9^.

**Figure 1.**
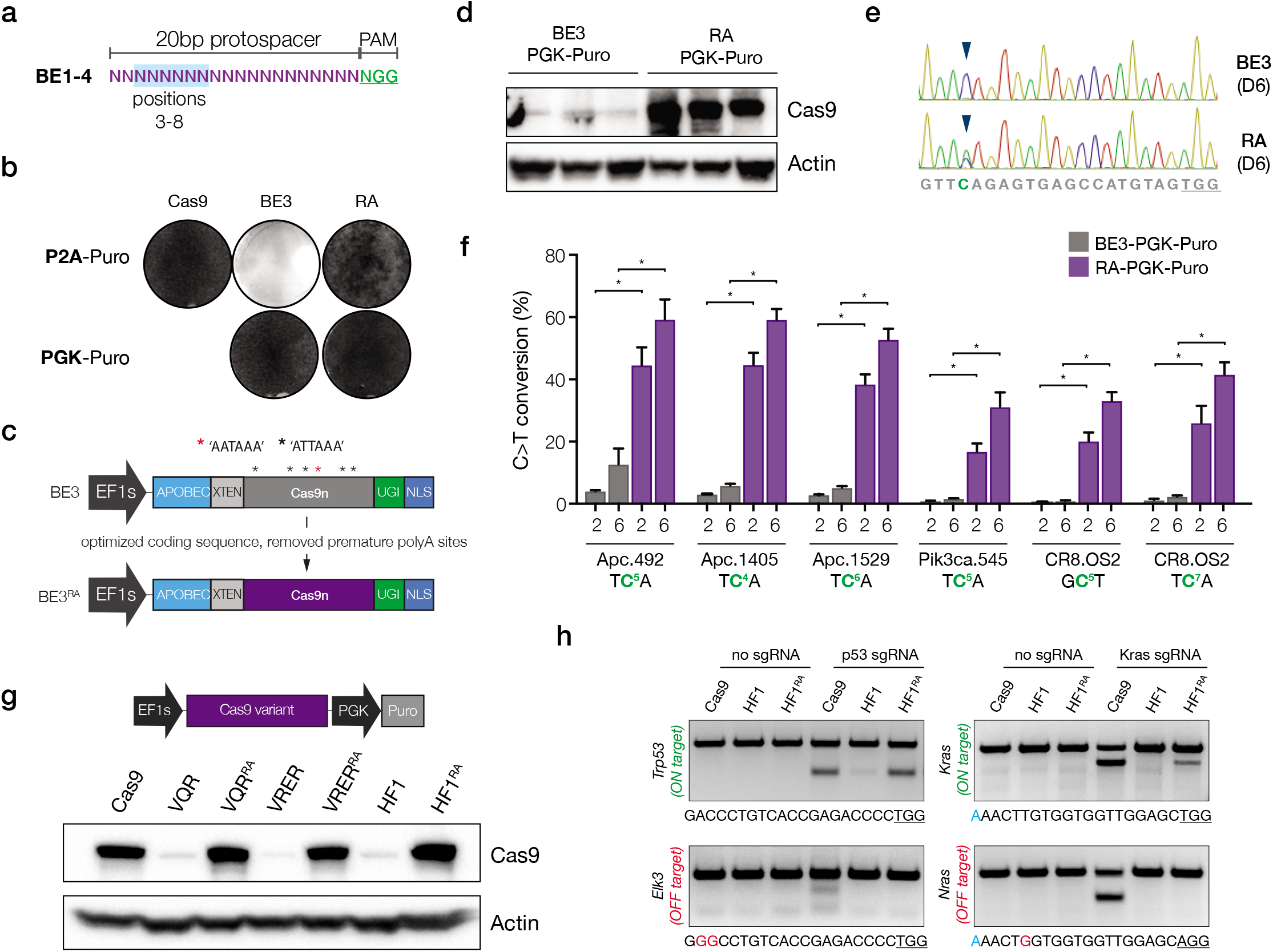
Optimizing the coding sequence of BE3 improves protein expression and target base editing. **a.** Schematic depiction of the canonical region of target base editing. Positions 3-8 (highlighted in blue) within the protospacer are susceptible to C>T conversion by BE3. The PAM is shown in green. **b.** Giemsa stained NIH/3T3 cells following transduction with indicated lentiviruses, and selection in puromycin for 6 days. **c.** Schematic representation of original BE3 (above), and codon optimized RA enzymes (below). **d.** Cas9 immunoblot of independently derived NIH/3T3 lines transduced with *BE3* or *RA* (n=3). **e.** Sanger sequencing chromatogram showing the target region of the Apc.1405 sgRNA. Arrow highlights cytosine at position 4 that shows dramatically increased editing by RA six days following sgRNA transduction. **f.** Frequency of target C>T editing across 5 different sgRNA targets, 2 and 6 days following sgRNA transduction, as indicated. Error bars represent +/-s.d., n = 3, asterisks (*) indicate a significant difference (p<0.05) between groups, using twoway ANOVA with Tukey’s correction for multiple testing. **g.** Western blot showing expression of original and optimized HF1 and PAM variant Cas9 proteins. **h.** T7 endonuclease assays on *Trp53* and *Kras* target sites, and off-target sites (*Elk3* and *Nras*, respectively) show that reassembled HF1 (HF1^RA^) improves on-target activity while maintaining little to no off-target cutting. Genomic target sites for each region are shown below. Of note, the slightly reduced on-target activity of HF1^RA^ at the Kras site may be due to the G-A mismatch at position 1 of the protospacer (highlighted blue).

We sought to use BE3 to engineer nonsense and missense mutations in cancer cell lines and intestinal organoids to study the impact of specific cancer-associated mutations. To do this we cloned a lentiviral vector in which BE3 was expressed from the EF1 short (EF1s) promoter and linked to a puromycin (puro) resistance gene via a P2A self-cleaving peptide (*pLenti-BE3-P2A-Puro*, BE3). Unexpectedly, and in contrast to *pLenti-Cas9-P2A-Puro* (Cas9), we were unable to generate puro-resistant cells, even in easily transduced cell lines (e.g. NIH/3T3s) (Figure 1b). We did not detect any significant difference in the production of viral particles, or integration of individual viral constructs into target cells (**Supplementary Figure 1**), suggesting the poor selection was due to low expression of the *BE3-P2A-Puro* cassette. To test this, we generated a new lentivirus in which the puro resistance (*Pac*) gene was driven by an independent (PGK) promoter (*pLenti-BE3-PGK-Puro*). This vector produced equivalent viral titer and target cell integration (**Supplementary Figure 1**), but, in contrast to the *BE3-P2A-Puro* vector, it enabled effective resistance to puro (Figure 1b, **Supplementary Figure 1c**).

Together, this work suggested an unresolved issue in the production of BE3 protein that was limiting effective base editing. Indeed, during cloning of the lentiviral constructs, we noted that the Cas9n DNA sequence in *BE3* was not optimized for expression in mammalian cells (Figure 1f), containing a large number of non-favored codons (**Supplementary Figure 2**) and 6 potential polyadenylation sites (AATAAA or ATTAAA) throughout the cDNA. We therefore reconstructed the BE3 enzyme using an optimized Cas9n sequence ^10^. The resulting reassembled *BE3* (*BE3^RA^*; hereafter “RA”) enabled efficient transduction and selection with either *PGK-Puro* or *P2A-Puro* lentivirus (Figure 1b, **Supplementary Figure 1**), markedly increased protein expression (Figure 1c), and most importantly, showed a 5-30 fold increase in target C>T conversion across multiple different genomic targets (Figure 1d,e; **Supplementary Figure 3**). Further, while C>T editing increased on average 15-fold, the level of unwanted insertions and deletions (indels), or undesired (C>A or C>G) editing remained low, indicating a significant improvement in the efficiency of the base editing over previous versions (**Supplementary Figure 4a,b**). Of note, we and others^11^ observed similar problems with expression of high-fidelity (*HF1*)^12^ and altered PAM specificity Cas9 variants^13^, which share the same Cas9 cDNA as BE3. In each case, this was corrected by re-engineering the construct with an optimized coding sequence (Figure 1g, **Supplementary Figure 5**)^11^. Importantly, the resulting increased expression of the HF1 enzyme (HF1^RA^) dramatically improved on-target DNA cleavage, while maintaining little or no off-target activity (Figure 1h).

**Figure 2.**
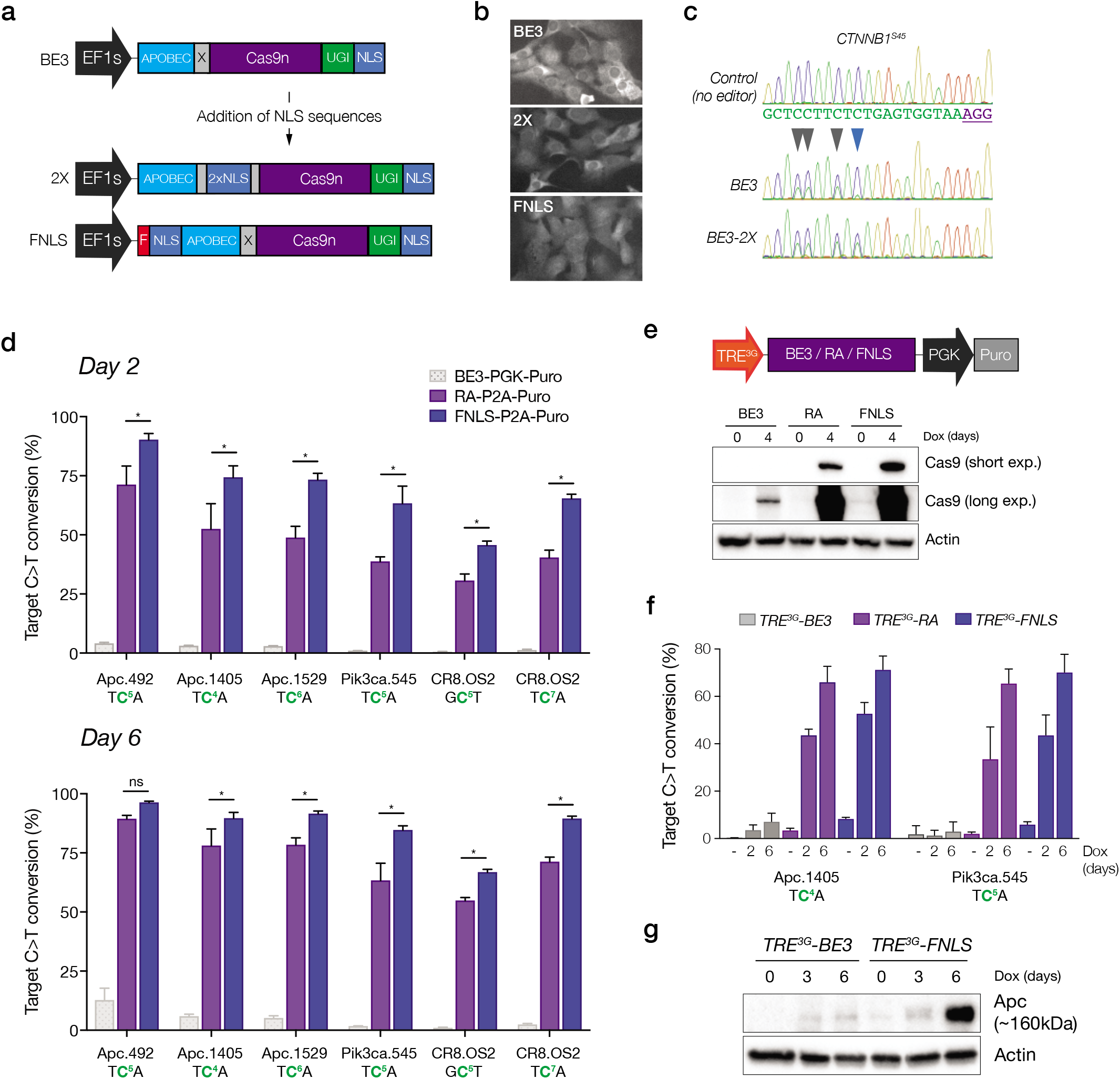
N-terminal NLS sequences increase the range and potency of target base editing. **a.** Schematic representation of original BE3 enzyme (above) and two new variants carrying NLS sequences within the XTEN linker (2X) or at the N-terminus (FNLS) **b.** Immunofluorescent staining of Cas9 in NIH/3T3 cells expressing RA, 2X, or FNLS. **c.** Sanger sequencing chromatogram showing increased editing of the cytosine at position 10 (blue arrow) within the protospacer of a *CTNNB1.S45* sgRNA. **d.** Frequency (%) of C>T conversion in NIH/3T3 cells transduced with *RA-* or *FNLS-P2A-Puro* lentiviral vectors 2 and 6 days following introduction of different sgRNAs, as indicated. Editing in *BE3-PGK-Puro* cells (from Figure 1e) is shown for comparison. Error bars represent s.e.m., n = 3, asterisks (*) indicate a significant difference (p<0.05) between groups, using a two-way ANOVA with Tukey’s correction for multiple testing. **f.** Schematic representation of dox-inducible BE3 lentiviral construct, and immunoblot of Cas9 in transduced and selected NIH/3T3 cells following treatment with or without dox (1μg/ml) for 4 days, as indicated. **g.** Frequency (%) of C>T conversion in NIH/3T3 cells transduced with *TRE^3G^-BE3, TRE^3G^-RA*, or *TRE^3G^-FNLS*, and sgRNA lentiviral vectors, 2 and 6 days following dox treatment. Error bars represent s.e.m., n = 3. **h.** Immunoblot showing induction of truncated (~160kDa) Apc product after target editing in NIH/3T3s expressing BE3 or FNLS.

**Figure 3.**
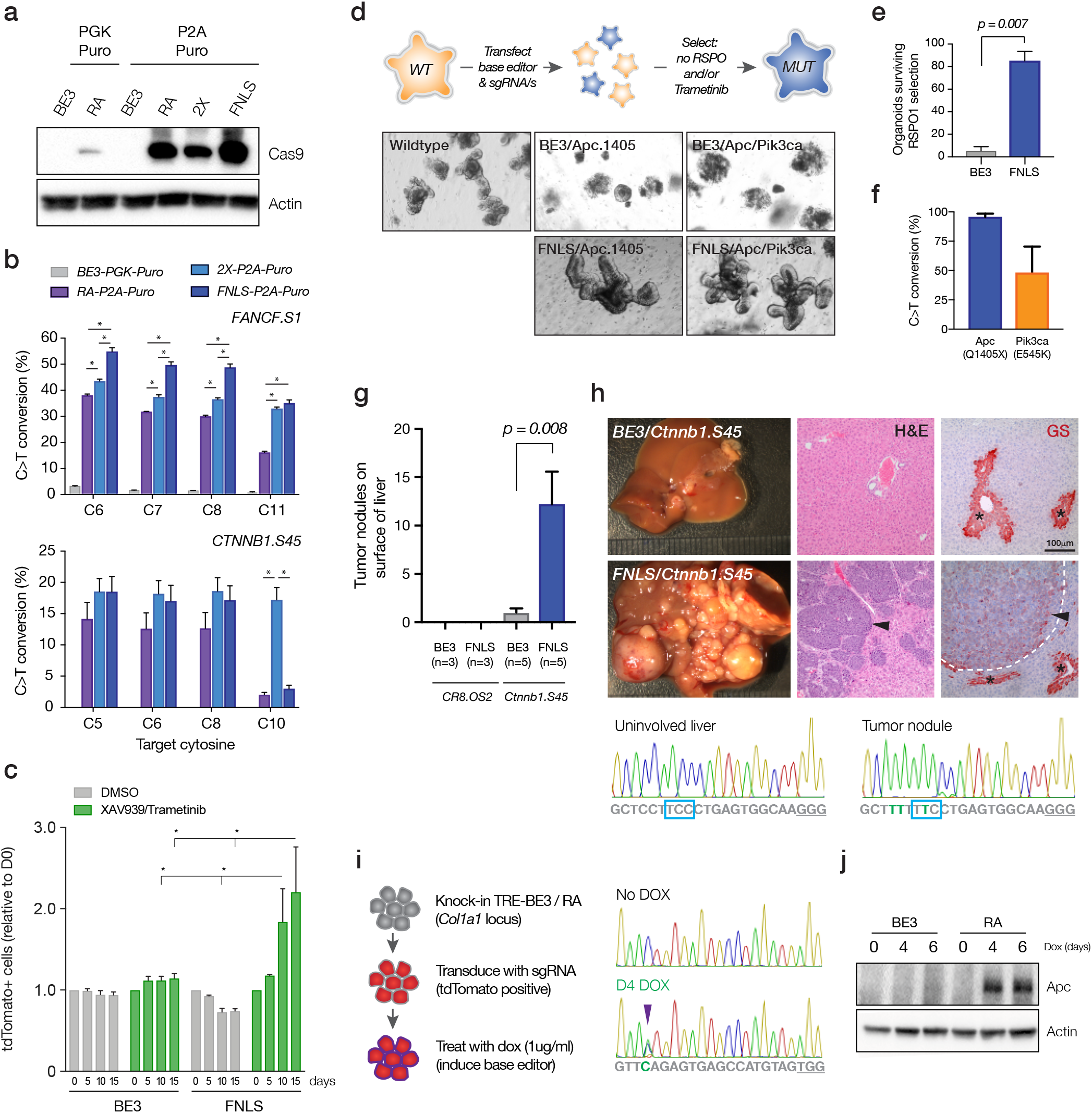
Optimized enzymes induce efficient base editing in a wide range of cell systems. **a.** Immunoblot of Cas9 in DLD1 cells transduced with base editor variants. **b.** Frequency (%) of C>T conversion in DLD1 cells transduced with base editor lentiviral vectors as indicated, analyzed 6 days following introduction of *FANCF.S1* and *CTNNB1.S45* sgRNAs. Error bars represent s.e.m., n = 3, asterisks (*) indicate a significant difference (p<0.05) between groups, using a two-way ANOVA with Tukey’s correction for multiple testing. Data from *BE3-PGK-Puro* is shown in place of *BE3-P2A-Puro*, as no editing was detected in cells transduced with the latter construct. **c.** Graph shows relative abundance of tdTomato-positive (sgRNA-expressing) cells in *BE3* and FNLS-transduced DLD1 cells, following treatment with DMSO, or XAV939 (1μM) and Trametinib (10nM). Bars in each case represent serial passages every 5 days, starting at day 0. Error bars represent s.e.m., n = 3, asterisks (*) indicate a significant difference (p<0.05) between groups, using a two-way ANOVA with Tukey’s correction for multiple testing. **d.** Schematic representation of the process of editing and selection in intestinal organoids. Images show wildtype mouse small intestinal organoids following editor/sgRNA transfection and selection by RSPO1 withdrawal (6 days). Only FNLS transfected organoids show consistent outgrowth of large budding organoids in the absence of RSPO1. Transfection with tandem sgRNAs targeting *Apc* and *Pik3ca* drives the generation of compound mutant organoids that survive RSPO1 withdrawal and treatment with 25nM Trametinib (see **Supplementary Figure 11**). **e.** Number of viable organoids six days following RSPO1 withdrawal. Error bars represent s.e.m., n = 2, two-sided t-test. **f.** Frequency of Apc Q1405X and Pik3ca E545K mutations in intestinal organoids following selection in RSPO1-free media. Error bars represent s.e.m., n = 3 independent transfections. **g.** Number of visible tumor nodules present of the liver of mice 4 weeks following hydrodynamic delivery of BE3 or FNLS, a mouse *Ctnnb1.S45* sgRNA and Sleeping Beauty transposon-based Myc cDNA. Error bars are s.e.m., n=3-5, as indicated; Statistical difference between groups calculated using a one-way ANOVA with Tukey’s correction for multiple testing. **h.** Representative images of tumor burden following editing of *Ctnnb1* with FNLS and BE3. *Right:* H&E and immunohistochemical staining for Glutamine synthetase (GS; red stain) of representative sections of livers from BE3 and FNLS transfected mice. Asterisks (*) highlight zone 1 regions of the liver that stain positively for GS. Arrows indicate tumors within the liver of FNLS-transfected mice. *Lower:* Sanger sequencing from uninvolved liver and a tumor nodule from an *FNLS/Ctnnb1.S45* sgRNA-transfected mice, showing near-complete editing of the *Ctnnb1* locus in tumor cells. Note: BE3 tumor nodules were too few and too small to dissect and obtain sequencing. **i.** *Left:* Schematic representation of the workflow to create inducible knock-in transgenic base editing allele, and measure editing following sgRNA transduction. *Right:* Sanger sequencing chromatogram shows editing of Apc in embryonic stem cells (ESCs) following 4 days of treatment with dox (1μg/ml). **j.** Immunoblot showing induction of the expected truncated allele of Apc in RA-expressing cells, but not in BE3 cells.

The addition of nuclear localization signal (NLS) sequences to the N-terminus of Cas9 can improve the efficiency of gene targeting ^14^ Indeed, despite the presence of a C-terminal NLS (Figure 2a), RA protein was largely excluded from the nucleus in transduced 3T3 cells (Figure 2b). We tested two different N-terminal positions for the NLS in case the inclusion of these sequences in one location interfered with APOBEC function: 1) with a FLAG epitope tag at the N-terminus (“*FNLS*”), and 2) within the XTEN linker that bridges APOBEC and Cas9n (*“2X”*) (Figure 2b). While 2X showed no obvious increase in nuclear targeting compared to RA, FNLS protein was more evenly distributed through the nucleus and cytoplasm of the cell (Figure 2b).

In transfection-based assays, FNLS increased editing across multiple target positions and sgRNAs (**Supplementary Figure 6**), while 2X significantly increased the range of editing toward cytosines at positions 10-11 in the protospacer (Figure 2c, **Supplementary Figure 6**). We next generated codon-optimized *2X-P2A-Puro* and *FNLS-P2A-Puro* lentiviral vectors and transduced mouse NIH/3T3 cells (**Supplementary Figure 7**). Two days following sgRNA transduction, FNLS-expressing cells showed greater than 50% C>T conversion for all sgRNAs tested (5/5), and by day six, 80-95% of all target cytosines were converted (Figure 2d). By contrast, only 1/5 sgRNAs showed >80% editing with RA (Figure 2d). The 2X construct showed a consistent trend toward increased editing, though in most cases this was not statistically significant (**Supplementary Figure 8**). Overall, FNLS increased target editing by 35% over RA, and up to 50-fold over the original BE3 construct (Figure 2d). FNLS expressing cells also produced fewer indels and non-desired (C>A and C>G) edits relative to RA (**Supplementary Figure 9**).

To provide a more flexible and controlled system for base editing, we generated lentiviral constructs whereby BE3, RA, or FNLS expression was controlled by a tetracycline responsive element (*TRE^3G^*) and doxycycline (dox)-dependent reverse tet-transactivator (rtTA3) (Figure 2e). As expected, we saw strong dox-mediated induction of RA and FNLS, but limited expression of the non-optimized BE3 construct (Figure 2f). Using sgRNAs targeting *Apc* and *Pik3ca*, we observed a time-dependent generation of target missense (*Pik3ca^E545K^*) and nonsense (*Apc^Q1405X^*) mutations (Figure 2g, **Supplementary Figure 10**). Consistent with earlier observations, both RA and FNLS dramatically increased editing efficiency relative to the original BE3 enzyme (Figure 2f), which for *Apc.1405*, translated to increased expression of a truncated Apc protein (Figure 2g).

Together, these data demonstrate that our re-engineered enzymes increase the range (2X) and efficiency (FNLS) of targeted base editing, and enable the effective use of dox-inducible systems to control the timing and duration of base editing. To demonstrate the utility and impact of the improved editors, we set out to engineer a series of precise genetic changes in four different model systems: human cancer cell lines, wildtype intestinal organoids, mouse embryonic stem cells, and mouse hepatocytes *in vivo*. In each case, we aimed to create missense mutations in *CTNNB1* (*S45F*), or nonsense mutations in *APC* (*Q1405X*), to drive hyperactivation of the WNT signaling pathway.

To first assess functional editing potential in human cancer cells, we generated DLD1 cells expressing each BE3 variant (Figure 3a). DLD1 cells are sensitive to combined inhibition of Tankyrase and MEK^15, 16^ but activating mutations in *CTNNB1* are predicted to induce resistance to this treatment ^17^. We introduced sgRNAs targeting codon (Serine) 45 of *CTNNB1*, or *FANCF.S1* as a neutral control (Figure 3b), and cultured the cells in Tankyrase (XAV939; 1uM) and MEK (Trametinib; 10nM) inhibitors (**Supplementary Figure 11**). At treatment initiation, 6 days following sgRNA transduction, RA, 2X and FNLS-expressing cells showed efficient editing (40-50%) at the FANCF target site in negative control cells, and *CTNNB1^S45F^* mutations at a frequency of 12-18% (Figure 3b). Most importantly, the editor-induced *CTNNB1^S45F^* mutations allowed bypass of WNT pathway suppression, and edited cells overtook non-edited cells in drug-treated cultures (Figure 3c, **Supplementary Figure 11d**). By contrast, BE3-expressing cells showed no detectable *CTNNB1* editing, and did not generate XAV939/Trametinib-resistant cultures (Figure 3c). Of note, as described in 3T3 cells (**Supplementary Figure 9**), FNLS significantly increased the ratio of desired (C>T) to non-desired (C>A, C>G) editing, and decreased the frequency of indels over the target site for both *CTNNB1.S45* and *FANCF.S1* (**Supplementary Figure 12**).

We next tested the ability of the optimized base editing tools to engineer cancer-associated mutations in wildtype intestinal organoids. Nonsense truncating *Apc* mutations are the most common genetic lesions observed in human colorectal cancers^18^, and drive WNT/RSPO ligand-independent activation of WNT signaling. To recreate this mutation, we co-transfected intestinal organoids with either BE3 or FNLS, and the *Apc.1405* sgRNA (Figure 3d). Strikingly, FNLS-transfected cultures showed a 10-fold increase in outgrowth of RSPO1-independent organoids, relative to BE3-transfected cells (Figure 3d,e). As expected, they carried a high frequency of targeted *Apc* disruption (>98%), and the vast majority of these alterations (>97%) were target C>T conversions, with fewer than 1% indels (Figure 3f). Further, the high editing efficiency of FNLS enabled multiplexed editing of independent genomic loci. Transfection of two tandem arrayed sgRNAs (*Apc.1405* and *Pik3ca.545*) produced *Apc*^Q1405X^/*Pik3ca^E545K^* mutant organoids (Figure 3f) that could survive and expand in the presence of a MEK inhibitor (Trametinib; 25nM) (**Supplementary Figure 13**), as has been described for HDR-generated *PIK3CA^E545K^* mutations in human intestinal organoids^19^. Thus, improved base editing tools provide a means to rapidly and precisely engineer multiple clinically relevant genetic lesions.

In human hepatocellular carcinoma (HCC), *CTNNB1* mutations are the primary mechanism of WNT-driven tumorigenesis in the liver. To explore the potential of base editors to drive tumor formation *in vivo*, we introduced BE3 or FNLS, a mouse *Ctnnb1.S45* sgRNA, and Myc cDNA to the livers of adult mice, via hydrodynamic transfection (Figure 3g). After 4 weeks, 3/5 BE3 transfected animals showed 1-2 small tumor nodules on the liver, while FNLS-transfected mice had dramatically increased disease burden with all mice (5/5) carrying multiple tumors (Figure 3g,h). Histologically, tumors resembled HCCs with trabecular and solid growth pattern, and showed upregulation of the β-catenin target, Glutamine synthetase ^20^ (GS; Figure 3h). As expected, tumor nodules showed efficient editing of the target *Ctnnb1* locus, to create activating S45F mutations (Figure 3h).

An alternate approach to *in vivo* somatic base editing is the generation of temporally regulated transgenic strains, which enables the manipulation of tissues and cell types that cannot be easily transfected *in vivo*, and avoids the potential immunogenicity of exogenous Cas9 delivery ^21,22^. We generated knock-in transgenic mouse embryonic stem cells (ESCs), in which *BE3* or *RA* expression is controlled by the TRE promoter. We chose *RA* for ESC targeting, as we noted low-level ‘leaky’ editing in 3T3s cells carrying *TRE^3G^-FNLS* lentivirus (Figure 2f). While *TRE-BE3* cells displayed no detectable editing following dox treatment, *TRE-RA* cells showed efficient conversion of target cytosines and generation of predicted mutant alleles (Figure 3i,j, **Supplementary Figure 14**). Together, these data show that optimized *RA* and *FNLS* constructs offer a flexible and efficient platform to engineer directed somatic alterations in animals.

Cas9-mediated base editing is a flexible and precise platform to engineer disease-relevant genetic alleles, and may provide a path forward for CRISPR-based therapeutics. Here, we developed optimized APOBEC-Cas9 fusion enzymes that dramatically improve base editing across multiple model systems and cell types, and offer a means for key for effective *in vitro* and *in vivo* base editing. Further, the generation of base editing enzymes that are efficient translated will be critically important for therapeutic approaches that rely on delivery of mRNA molecules ^23^. Together, we believe these optimized enzymes make base editing technology a feasible and accessible option for a wide range of research and therapeutic applications.

## Acknowledgements

This work was supported by a project grant from the NIH/NCI (CA195787-01), a U54 grant from the NIH/NCI (U54OD020355), project grant from the Starr Cancer Consortium (I10-0095), a Research Scholar Award from the American Cancer Society (RSG-17-202-01), and a Stand Up to Cancer Colorectal Cancer Dream Team Translational Research Grant (SU2C-AACR-DT22-17). Stand Up to Cancer is a program of the Entertainment Industry Foundation. Research grants are administered by the American Association for Cancer Research, a scientific partner of SU2C. MPZ was supported in part by National Cancer Institute (NCI) Grant NIH T32 CA203702. EMS was supported by a Medical Scientist Training Program grant from the National Institute of General Medical Sciences of the National Institutes of Health under award number T32GM07739 to the Weill Cornell / Rockefeller / Sloan-Kettering Tri-Institutional MD-PhD Program, and an F31 Award from the NCI/NIH under grant number 1 F31 CA224800-01. FJSR was supported by the MSKCC TROT program (5T32CA160001) and is an HHMI Hanna Gray Fellow. SWL is the Geoffrey Beene Chair of Cancer Biology and an Investigator of the Howard Hughes Medical Institute. DFT is supported by the Helmholtz Association (VH-NG-1114) and by the German Research Foundation (DFG) project B05, SFB/TR 209 “Liver Cancer”. LED was supported by a K22 Career Development Award from the NCI/NIH (CA 181280-01). The content is solely the responsibility of the authors and does not necessarily represent the official views of the NIH.

## Author Contributions

MPZ and EMS performed experiments, analyzed data and wrote the paper. AK, MF, AS, EM, TH, JT, and FSR performed experiments and analyzed data. MB and AYS performed and analyzed in vivo experiments. DFT designed and supervised in vivo experiments. JS, SWL, and CV supplied critical unpublished reagents. LED performed and supervised experiments, analyzed data, and wrote the paper.

## Methods

### Cloning

All primers, Ultramers, and gBlocks used for cloning are listed in **Supplementary Table 2**. pCMV-BE3-2X (CMV-2X) and pCMV-BE3-FNLS were generated by Gibson assembly, combining an XmaI-digested (2X) or NotI-digested (FNLS) pCMV-BE3 backbone with DNA Ultramers (BE3-2X NLS or T7-FLAG-NLS). Double stranded DNA from Ultramers was generated by PCR amplification using primers XTEN-NLS_F/XTEN-NLS_R, and T7-FLAG_F/T7-FLAG_R primers, respectively. *pLenti-BE3-PGK-Puro* (*LBPP*) was generated by Gibson assembly combining 4 DNA fragments: i) PCR amplified EF1s promoter (FSR-19/FSR-20), ii) PCR amplified BE3 cDNA (FSR-114/FSR-115), iii) PCR amplified PGK-Puro cassette (FSR-16/FSR-17), and iv) a BsrGI/PmeI digested pLL3-based lentiviral backbone. *pLenti-BE3^R^-PGK-Puro (LRPP)* was generated by Gibson assembly, combining a PCR amplified BE3RA cDNA (BE3RA-PGKPuro_F/BE3RA-PGKPuro_R) and an NheI/AvrII digested BE3-PGK-Puro backbone. *pLenti-FNLS-PGK-Puro (LFPP)* was generated by restriction cloning a FLAG-NLS-APOBEC BamHI(blunt)/EcoRI-digested fragment, into an NheI(blunt)/EcoRI-digested pLenti-BE3RA-PGK-Puro backbone. *pLenti-BE3^m^-P2A-Puro (LR2P)* was generated by Gibson assembly, combining 4 DNA fragments: i) a PCR amplified APOBEC-XTEN cDNA (BE3RA_APOBEC_F /BE3RA_XTEN_R), ii) PCR amplified Cas9n (BE3RA_Cas9n_F/BE3RA_Cas9n_R), iii) PCR amplified UGI (BE3RA_UGI_F/BE3RA_UGI_R), and a BamHI/NheI digested *pLenti-Cas9-P2A-Puro* viral backbone. Note: some wobble positions were altered within the UGI (SGGS) linker to avoid complications during Gibson assembly because of an identical region downstream of UGI. *pLenti-FNLS-P2A-Puro (LF2P)* was generated by restriction cloning a PCR amplified (BamHI-FLAG_F/APOBEC-RI_R) BamHI/EcoRI-digested, FLAG-NLS-APOBEC fragment into a BamHI/EcoRI digested pLenti-BE3RA-P2A-Puro backbone. *pLenti-2X-P2A-Puro (LX2P)* was generated by Gibson assembly, combining a PCR-amplified APOBEC-2XNLS fragment (BE3RA_APOBEC_F/BE3RA_XTEN_R) and an BamHI/XmaI digested pLenti-BE3RA-P2A-Puro backbone. *pLenti-TRE^3G^-BE3-PGK-Puro (L3BP)* was generated by Gibson assembly, combining a PCR-amplified TRE3G promoter (3G_F/3G_R), and APOBEC fragment (APOBEC_F/BE3RA_XTEN_R) with an XmaI digested pLenti-BE3-PGK-Puro backbone. *pLenti-TRE^3G^-BE3^RA^-PGK-Puro (L3RP)* was generated by Gibson assembly, combining a PCR-amplified TRE3G promoter (3G_F/3G_R), and APOBEC fragments (APOBEC_F/BE3RA_XTEN_R) with an XmaI digested pLenti-BE3RA-PGK-Puro backbone. *pLenti-TRE^3G^-FNLS-PGK-Puro (L3FP)* was generated by Gibson assembly, combining a PCR-amplified TRE3G promoter (3G_F/3G_R), and FNLS-APOBEC fragments (FNLS-APOBEC_F/BE3RA_XTEN_R) with an XmaI digested pLenti-BE3RA-PGK-Puro backbone. *pCol1a1-TRE-BE3 (cTBE3)* was generated by Gibson assembly, combining a PCR-amplified BE3 cDNA (cTRE_BE3_F/cTRE_BE3_R) with an EcoRI-digested pCol1a1-TRE backbone. *pCol1a1-TRE-BE3RA (cTBE3™)* was generated by a two-step strategy. First, using Gibson assembly to introduce a PCR amplified UGI fragment (UGI_F/UGI_R) into a XhoI-digested pCol1a1-TRE-Cas9n backbone (*Col1a1-TRE-Cas9n-UGI*). Second, by restriction cloning a PCR-amplified, XhoI/EcoRV-digested APOBEC-XTEN-Cas9n (APOBEC_F2/APOBEC_R2) fragment into an EcoRV-digested Col1a1-TRE-Cas9n-UGI backbone. *pLenti-U6-sgRNA-tdTomato-P2A-Blas (LRT2B)* was generated by Gibson assembly, combining a PCR-amplified EFs-TdTomato-P2A-Blasticidin fragment (pLRT2B_EFs_F/pLRT2B_WPRE_R) with an XhoI/BsrGI-digested pLenti-U6-sgRNA-GFP (LRG) backbone. *pLenti-VQR-P2A-Puro (LQ2P), pLenti-VRER-P2A-Puro (LER2P)*, and *pLenti-HF1-P2A-Puro (LH2P)* were generated by Gibson assembly, combining PCR amplified Cas9 variants (from Addgene stocks: #65771, #65773, and #72247, respectively; primers: KJ_Cas9_F/KJ_Cas9_R) with BamHI/NheI-digested pLenti-P2A-Puro backbone. *pLenti-VQR^m^-P2A-Puro (LQR2P), pLenti-VRER^R4^-P2A-Puro (LERR2P)*, and *pLenti-HF1^R4^-P2A-Puro (LHR2P)* were generated by Gibson assembly, combining one of two PCR amplified regions of the 3’ half of Cas9 (Cas9_RA_5F/Cas9_RA_5R or Cas9_RA_3F/Cas9_RA_3R), with gBlock fragments containing appropriate point mutations (VQR_GB, VRER_GB, or HF1_GB), and an EcoRV/NheI-digested *pLenti-Cas9-P2A-Puro* backbone. All constructs described above are schematized in **Supplementary Figure 13**.

### Cell culture, transfection, and transduction

#### Culture

HEK293T (ATCC CRL-3216) and DLD1 cells (ATCC CCL-221) were maintained in Dulbecco’s Modified Eagle’s Medium (Corning) supplemented with 10% (v/v) fetal bovine serum (FBS), at 37° with 5% CO_2_. NIH/3T3 (ATCC CRL-1658) were maintained in Dulbecco’s Modified Eagle’s Medium (Corning) supplemented with 10% (v/v) bovine calf serum (CALF). Mouse KH2 embryonic stem cells were maintained on irradiated MEF feeders in M15 media containing LIF, as previously outlined ^24^

#### Transfection

For transfection-based editing experiments in HEK293Ts, cells were seeded on a 12-well plate at 80% confluence and co-transfected with 750ng of base editor (BE), 750ng of sgRNA expression plasmid, and 4.5μl of polyethyleminine (PEI; 1mg/ml). Cells were harvested for genomic DNA, 3 days post-transfection. For virus production, HEK293T cells were plated in a 6 well-plate and transfected 12 hours later (95% confluence) with a prepared mix in DMEM media (no supplements) containing 2.5μg of lentiviral backbone, 1.25μg of PAX2, 1.25μg of VSV-G, and 15μl of PEI (1mg/ml). 36hrs following transfection, media was replaced with target cell collection media and supernatants were harvested every 8-12hrs up to 72hrs post transfection. ESCs collal-targeting constructs were introduced via nucleofection in 16-well strips, using buffer P3 (Lonza Inc., V4XP-3032) in a 4D Nucleofector with X-unit attachment (Lonza Inc.). Two days following nucleofection, cells were treated with media containing 150ug/ml Hygromycin B and individual surviving clones were picked after 9-10 days of selection. Two days after clones were picked Hygromycin was removed from the media and cells were cultured in M15 thereafter. To confirm integration at the *collal* locus we used a multiplex *collal* PCR ^24^.

#### Transduction

7.5 × 10^4^ NIH/3T3 or DLD1 cells were plated on 6-well plate. 24hrs following plating, cells were transduced with viral supernatants in the presence of polybrene (8μg/μl). Two days after transduction cells were selected in Puromycin (2 ug/ml), Blasticidin S (4 μg/ml) or G418 (Neomycin; 500 ug/ml). 500k ESCs were plated in 6-well plates on gelatin and spinocculated (90 mins, 32°C, 2100 rpm) with 150 μl of concentrated lentiviral particles (using 100mg/ml polyethylene glycol, Sigma Aldrich P4338) in 1 ml of media containing polybrene (8μg/μl). After spin, media was replaced.

#### Fluorescent competitive assays

DLD1 cells expressing BE3, RA, 2X, or FNLS were transduced with LRT2B-CTNNB1^S45^ or LRT2B-FANCF^S1^, selected with Blasticidin for 4 days and mixed at defined proportions with parental cells. 5 × 10^4^ mixed cells were seeded in 96 well plates, treated with DMSO or 1μM XAV939 + 10nM Trametinib every 48h and remaining tdTomato-positive cells were tracked every 5 days by flow cytometry using a BD-Accuri C6 cytometer.

### Organoid isolation, culture, transfection, and transduction

Organoid isolation was performed as previously described ^25, 26^. Briefly, 15 cm of the proximal small intestine was removed, flushed and washed with cold PBS. The intestine was then cut into 5 mm pieces and placed into 10 ml cold 5mM EDTA-PBS and vigorously resuspended using a 10ml pipette. The supernatant was aspirated and replaced with 10ml EDTA and placed at 4ºC on a benchtop roller for 10 minutes. This was then repeated for a second time for 30 minutes. The supernatant was aspirated and then 10ml of cold PBS was added to the intestine and resuspended with a 10ml pipette. After collecting this 10ml fraction of PBS containing crypts, this was repeated and each successive fraction was collected and examined underneath the microscope for the presence of intact intestinal crypts and lack of villi. The 10ml fraction was then mixed with 10ml DMEM Basal Media (Advanced DMEM F/12 containing Pen/Strep, Glutamine, 1mM N-Acetylcysteine (Sigma Aldrich A9165-SG)) containing 10 U/ml DNAse I (Roche, 04716728001), and filtered through a 100μm filter. It was then filtered through a 70μm filter into an FBS (1ml) coated tube and spun at 1200 RPM for 3 minutes. The supernatant was aspirated and the cell pellet (purified crypts) were resuspended in basal media, mixed 1:10 with Growth Factor Reduced Matrigel (BD, 354230), and plated in multiple wells of a 48 well plate. After polymerization for 15 mins at 37C, 250μl of small intestinal organoid growth media (Basal Media containing 50 ng/mL EGF (Invitrogen PMG8043), 100ng/ml Noggin (Peprotech 250-38), and R-spondin (conditioned media) was then laid on top of the Matrigel.

#### Maintenance

Media was changed on organoids every two days and they were passaged 1:4 every 5-7 days. To passage, the growth media was removed and the Matrigel was resuspended in cold PBS and transferred to a 15ml falcon tube. The organoids were mechanically disassociated using a p1000 or a p200 pipette and pipetting 50-100 times. 7 ml of cold PBS was added to the tube and pipetted 20 times to fully wash the cells. The cells were then centrifuged at 1000 RPM for 5 minutes and the supernatant was aspirated. They were then resuspended in GFR Matrigel and replated as above. For freezing, after spinning the cells were resuspended in Basal Media containing 10% FBS and 10% DMSO and stored in liquid nitrogen indefinitely.

#### Transfection

Murine small intestinal organoids were cultured in medium containing CHIR99021 (5μM) and Y-27632 (10μM) for 2 days prior to transfection. Cells suspensions were produced by dissociating organoids with TrypLE™ express (Invitrogen #12604) for 5 min at 37°C. After trypsinization, cell clusters in 300μl transfection medium were combined with 100μl DMEM/F12-Lipofectamine2000 (Invitrogen #11668)-DNA mixture (97ul-2ul-1ug), and transferred into a 48-well culture plate. The plate was centrifuged at 600g at 32°C for 60 min, followed by another 6h incubation at 37°C. The cell clusters were spun down and plated in Matrigel. For selecting organoids with *Apc* mutations, exogenous RSPO1 was withdrawn 2-3 days after transfection. For selection of Pik3ca alterations, organoids were cultured in medium containing Trametinib (25nM) for 1 week.

#### Transduction

Organoids were prepared as described for transfection to generate small cell clusters. They were then mixed with viral supernatant and polybrene (8μg/μl) before spinocculation and incubation at 37C. Cell clusters were plated as described for transfection and, after 48 hours, organoids were selected with Puromycin (2μg/μl) for 5-7 days (including at least one passage).

### Hydrodynamic delivery

All animal experiments were authorized by the regional board Karlsruhe, Germany (animal permit number G178/16) or the Institutional Animal Care and Use Committee (IACUC) at Weill Cornell Medicine (2014-0038). Eight week-old C57Bl6/N mice (Charles River, Sulzfeld, Germany) were injected with 0.9% sterile sodium-chloride solution containing 20μg *pLenti-BE3-P2A-Puro* or *pLenti-FNLS-P2A-Puro*, 10μg of respective sgRNA vector, and 5μg pT3 EF1a-myc, as well as 1μg CMV-SB13. The total injection volume corresponded to 20% of the individual mouse body weight and was injected into the lateral tail vein in 5-7 seconds. No animals were excluded from the analyses; the investigators were not blinded for the analyses.

### Lentiviral titer assay

Lentiviral titers were calculated using a quantitative PCR-based kit (LV900; Applied Biological Materials Inc.), according to the manufacturer’s instructions. Briefly, 2μl of unconcentrated viral supernatant was lysed for 3mins at room temperature, and the crude lysis was used to perform qPCR amplification. The concentration of viral particles was calculated as described in the protocol (http://www.abmgood.com/High-Titer-Lentivirus-Calculation.html).

**Genomic DNA isolation.** Cells were lysed in genomic lysis buffer (10 mM Tris pH 7.5, 10 mM EDTA, 0.5% SDS, 400 μg/ml Proteinase K) for at least 2hrs at 55C. Following Proteinase K heat inactivation at 95C for 15 mins, 0.5 volumes of 5M NaCl was added and centrifuged for 10mins at 15K rpm. Supernatant was mixed with 1 volume of isopropanol and DNA precipitates were washed in EtOH 70% before resuspension in 10 mM Tris pH 8.0.

### Puro copy number assay

For quantification of lentiviral integrations in transduced cells we used a custom-designed Taqman copy number assay (Invitrogen) to detect the *Pac (puroR*) gene. Amplification was conducted on QuantStudio 6 Real-Time PCR system (Applied biosystems), using Taqman master mix reagent (Applied biosystems) and specific primers and probe (forward-5’GCGGTGTTCGCCGAGAT; reverse-5’GAGGCCTTCCATCTGTTGCT; probe (FAM) CCGGGAACCGCTCAACTC)

### Protein analysis

293Ts and 3T3s were scraped from a confluent well of a 6 well plate in 100ul RIPA buffer then centrifuged at 4°C at 13,000rpm to collect protein lysates. DLD1 cells were pelleted from a confluent well of a 6 well plate at 1000rpm x 4 min, resuspended in 200 ul RIPA Buffer, then centrifuged at 4°C at 13,000rpm to collect protein lysates. Organoids were collected from confluent well of a 12 well plate (~100 ul Matrigel) in 200 ul Cell Recovery Solution (Corning, #354253), incubated on ice for 20 min, then pelleted at 300 g x 5 min. The pellet was then resuspended in 20 ul RIPA buffer, and centrifuged at 4°C at 13,000rpm to collect protein lysates. ESCs were collected at the indicated time points and filtered through a 40μm cell strainer (Fisher Scientific) to remove feeders, then pelleted at 1000 rpm x 4 min and resuspended in 100 μl RIPA Buffer. Samples were centrifuged at 4°C at 13,000rpm to collect protein lysates. Antibodies used for western blot analyses were: Cas9 (BioLegend, #844301), Actin (Abcam, #ab49900), and Apc (Millipore, MABC202).

### IF staining & microscopy

2 x10^4^ editor-expressing 3T3s were plated in a chamber slide. 24 h after cells were wash in PBS and fix in PBS-4% PFA solution 20 min at RT and incubated in permeabilization buffer (PBS-0.5% Triton X-100) for 10 min on ice. Then, cells were stained with Cas9 (BioLegend, San Diego, CA, USA, #844301) antibody at 4°C overnight. A donkey anti-mouse Alexa 594 (Thermo Fisher Scientific, Waltham, MA, USA, # A21203) was used as secondary antibody.

### Immunohistochemistry

Slides containing 3μm thick liver sections were deparaffinized and rehydrated using a descending alcohol series. For antigen retrieval slides were cooked in sodium-citrate buffer (pH 6.0) in a pressure cooker for 8 minutes. Subsequently, endogenous HRP was blocked for 10 minutes in 3% H_2_O_2_. Slides were blocked with in PBS containing 5% BSA for one hour before incubation with the primary antibody (Anti-mouse Glutamine Synthetase, BD610517, BD, Heidelberg Germany) overnight (1:200 dilution in PBS/5% BSA). Slides were washed three times and staining was visualized using DAKO Real Detection System (K5003, DAKO, Hamburg, Germany) according to the manufacturers instructions.

### PCR amplification for MiSeq

Target genomic regions of interest were amplified by PCR using primers pairs listed in Supplementary oligo table. PCR was performed with Herculase II Fusion DNA polymerase (Agilent Technologies, Palo Alto, CA, USA, #600675) according to the manufacturer’s instructions using 200 ng of genomic DNA as a template and under the following PCR conditions: 95°C x 2 min, 95 °C - 0:20 à 58 °C - 0:20à 72 °C - 0:30 × 34 cycles, 72 × 3 min. PCR products were column purified (Qiagen) for analysis by Sanger sequencing or MiSeq.

### DNA Library Preparation and MiSeq

DNA library preparations and sequencing reactions were conducted at GENEWIZ, Inc. (South Plainfield, NJ, USA). NEB NextUltra DNA Library Preparation kit was used following the manufacturer’s recommendations (Illumina, San Diego, CA, USA). Adapter-ligated DNA was indexed and enriched by limited cycle PCR. The DNA library was validated using TapeStation (Agilent Technologies, Palo Alto, CA, USA), and was quantified using Qubit 2.0 Fluorometer.

The DNA library was quantified by real time PCR (Applied Biosystems, Carlsbad, CA, USA). The DNA library was loaded on an Illumina MiSeq instrument according to manufacturer’s instructions (Illumina, San Diego, CA, USA). Sequencing was performed using a 2×150 paired-end (PE) configuration. Image analysis and base calling was conducted by the MiSeq Control Software (MCS) on the MiSeq instrument, and verified independently using a custom workflow in Geneious R11.

